# Cryo-EM structure of alpha-synuclein fibrils amplified by PMCA from PD and MSA patient brains

**DOI:** 10.1101/2021.07.08.451588

**Authors:** Domenic Burger, Alexis Fenyi, Luc Bousset, Henning Stahlberg, Ronald Melki

**Author notes:** Contributed equally (Domenic Burger, Alexis Fenyi). Correspondence to: Luc Bousset, Henning Stahlberg and Ronald Melki.

## Abstract

Synucleinopathies are neurodegenerative diseases related to the aggregation of the protein alpha-synuclein (aSyn). Among these diseases, Parkinson’s disease (PD) and multiple system atrophy (MSA) are most prevalent. aSyn can readily form different fibrillar polymorphs, if exposed to an air-water interface or by templating with pre-existing fibrils. We here report the structures of three fibrillar polymorphs that were obtained after seeding monomeric aSyn with PD and MSA patients brain homogenates using protein misfolding cyclic amplification (PMCA). Seeding with a control brain homogenate did not produce fibrils, and seeding with other *in vitro* generated fibrillar polymorphs as a control faithfully produced polymorphs of a different type. The here determined fibril structures from PD and MSA brain tissue represent new folds, which partly resemble that of previously reported *in vitro* generated fibrils from Y39 phosphorylated aSyn protein. The relevance of these fibrils for synucleinopathies in humans remains to be further investigated.

**Impact Statement:** Neurodegenerative diseases such as Parkinson’s disease (PD) and Multiple System Atrophy (MSA) are suspected to be causatively related to the prion-like propagation of aggregates of the protein alpha-synuclein (aSyn). The fibril structures reported here were obtained after seeding from diseased human brain homogenate and differ from all previously published aSyn fibril arrangements. In case these fibrils would turn out to be the long sought causative agents of these diseases, their structures might lead to the development of therapeutic strategies to modify these diseases and to a better understanding of the mechanistic processes that lead to neurodegeneration and spreading of the diseases.

## Introduction

Synucleinopathies are neurodegenerative disorders that are related to the dysfunction of aSyn and lead to consequential damage in neuronal cells with eventual cell death. These include PD, MSA, dementia with Lewy bodies, and others. aSyn-rich cellular inclusions, termed Lewy bodies (LB) in cell bodies and Lewy neurites (LN) in neurons, and cytoplasmic inclusions in oligodendrocytes, are central hallmarks found in the tissue of *post-mortem* brain. This commonality suggests a shared pathological pathway (Jellinger, 2009; Martí et al., 2003) within synucleinopathies and other types of neurodegeneration. aSyn is a 140 amino acid long cytosolic protein that is proposed to be involved in numerous cellular functions with the detailed mechanisms still subject to debate (Bellani et al., 2010; Burré et al., 2010; Emamzadeh, 2016; Fusco et al., 2016; Huang et al., 2019; Villar-Piqué et al., 2016). The protein is highly dynamic and acts like a protein-chameleon, as it can adopt different conformations in different environments and purposes (Munishkina et al., 2003; Uversky, 2003). Natively, aSyn is an unstructured monomeric protein. Upon interaction with acidic phospholipids, seven imperfect amino-terminal repeats, consisting of 11 amino acid long stretches centered on the consensus KTKEGV, adopt an alpha-helical conformation (Bartels et al., 2011; Chandra et al., 2003; Ulmer et al., 2005). The lysine rich N-terminal domain (residues 1-60) is involved in the lipid binding function of aSyn. It is followed by a hydrophobic sequence, termed the non-amyloid-beta component (NAC, residues 61-95). The NAC has been associated with a role in the pathological aggregation of aSyn (Beyer, 2006; Goedert, 2001). The remaining unstructured acidic carboxy-terminal domain (residues 96-140) can be subject to truncations which may promote fibril formation as well (Crowther et al., 1998; Li et al., 2005).

Full-length or truncated aSyn can aggregate or incorporate into amyloid fibrils, in which the protein adopts a beta strand-rich conformation, by either primary nucleation or templating events of pre-existing fibrils. In these fibrils, aSyn monomers form hydrogen bonds between their carbon backbone and that of the neighboring aSyn monomers, thereby forming tightly packed repetitive stacks that are often composed of two protofibrils. The spacing between monomers along the fibril axis is independent of the protein side chains and defined by the hydrogen bond length, giving a repetition of around 4.7Å. We previously showed that different experimental conditions can lead to the formation of different beta-strand rich secondary structures, and consequently lead to different fibril strains (Bousset et al., 2013; Verasdonck et al., 2016). These amyloids differ in (i) the structure that the aSyn molecules adopt within the fibrillar scaffold (i.e., the fibril strain), and (ii) in the surfaces that the fibrils expose to the solvent, which defines their interaction with partner molecules or cellular processes (Shrivastava et al., 2017).

Evidence supporting the propagation of aSyn-rich inclusions in a prion-like manner within the nervous system originates from two fundamental observations. The first comes from immunohistochemical studies which showed that Lewy pathology is present not only in different brain areas, but also in peripheral nerves. As the affected tissues are connected by neural pathways a “causative agent”, initially affecting the olfactory bulb and/or the autonomic nervous system innervating the gut, was proposed to spread progressively through those connections. This phenomenon is considered to be the origin of disease progression, as described by the neuropathological staging proposed by Braak and coworkers (Braak et al., 2006, 2003; Del Tredici and Braak, 2016). The second evidence supporting the spread of aSyn-rich inclusions comes from patients who were developing synucleinopathies and who had received grafts of fetal neurons into their brains years before their death. Upon autopsy of those patient brains, the grafted neurons were found to be affected by Lewy pathology (Kordower et al., 2008; Li et al., 2008). It appeared as if aSyn-rich deposits had moved from affected cells in the host brain to the naïve neurons within the grafts, where they triggered the further aggregation of endogenous aSyn. The evidences of spreading and amplification of pathology led to the design of numerous cell culture and *in vivo* studies, supporting the view that aSyn fibrils made in test tubes and aSyn-rich deposits purified from model animals or patients have prion-like properties (Angot et al., 2012; Aulić et al., 2014; Bousset et al., 2013; Brahic et al., 2016; Danzer et al., 2009; Desplats et al., 2009; Freundt et al., 2012; Hansen et al., 2011; Holmqvist et al., 2014; Kordower et al., 2011; Lee et al., 2010; Luk et al., 2012a, 2012b, 2009; Masuda-Suzukake et al., 2013; Mougenot et al., 2012; Peelaerts et al., 2015; Rey et al., 2016, 2013; Reyes et al., 2015; Sacino et al., 2014, 2013; Shimozawa et al., 2017; Tran et al., 2014; Volpicelli-Daley et al., 2011; Zhang et al., 2018). Those studies have established that besides their ability to elongate by recruitment of monomeric aSyn molecules, exogenous aSyn fibrillar assemblies possess templating capacity both *in vitro* and *in vivo* (Bousset et al., 2013; Peelaerts et al., 2015). Similarly, aSyn-rich inclusions purified from patient brains affected by different synucleinopathies as well as whole brain homogenates from such patients, but not from control cases, have been demonstrated to promote monomeric aSyn assembly into fibrillar structures that exhibit distinct shapes, proteolytic profiles and pathogenic properties *in vivo* indicating a templating effect of the pathogenic fibril seeds (Candelise et al., 2019; Fenyi et al., 2019; Prusiner et al., 2015; Shahnawaz et al., 2020; Strohäker et al., 2019; Van der Perren et al., 2020; Watts et al., 2013).

These findings show that a detailed understanding of the atomic arrangement of aSyn within structurally and pathologically distinct fibrillar assemblies is important in order to interfere with their capacity to elongate and interact with partner molecules within neuronal cells. Furthermore, high-resolution structural information is required to identify the part of the protein responsible for fibril templating as well as the surfaces of pathogenic aSyn fibrils exposed to the solvent. Together these data help to define how aSyn fibrils interact with partner molecules, cellular compartments or other native aSyn monomers, causing loss of protein function, interference with intracellular processes, fibril elongation and seeding, and cell-to-cell propagation. Several structures of aSyn fibrils that were assembled de novo from wildtype, truncated, mutant and/or post-translationally modified aSyn protein have been solved (Guerrero-Ferreira et al., 2020, 2019, 2018; B. Li et al., 2018; Y. Li et al., 2018). Those studies have confirmed the ability of aSyn to assemble into a wide range of structurally heterogeneous fibrillar assemblies.

As aSyn can be found in *post-mortem* brain tissue in LB and LN inclusions (Spillantini et al., 1998), attempts to generate *in vitro* fibrils have been made (Crowther et al., 1998). For amyloid-β, it was shown that an air-water interface influences the secondary structure motif and favors conversion of alpha-helical domains into beta-sheet rich assemblies (Schladitz et al., 1999). Also, aSyn fibrils form spontaneously within a few hours, if purified monomeric aSyn is kept in solution in the presence of an air-water interface, a process that is accelerated if the sample is at room or higher temperatures (Campioni et al., 2014; Pronchik et al., 2010). Storing the sample in a conventional Eppendorf tube with air above the solution, for example, fulfills such conditions and triggers spontaneous *in vitro* fibril formation. Instead, if monomeric aSyn is kept in solution in the absence of an air-water interface, as for example when kept within a dialysis bag without air bubbles and immersed in buffer solution, then in the absence of fibril seeds no fibril formation was observed in our hands even after weeks of maintaining the solution at room temperature.

Despite the ability of *in vitro* generated fibrils to trigger distinct pathology in model animals, the recent structural analysis of aSyn fibrillar assemblies purified from MSA patient brains by the groups of Goedert and Scheres revealed aSyn fibrils of entirely different structures than that of all previously studied *in vitro* generated fibrils (Schweighauser et al., 2020). The purification protocol used in that study involved solubilization of brain membranes by sarkosyl, which may or may not have affected the state of the fibrils.

It is important to understand if and how specific fibrillar assemblies impact intracellular processes and cause neurodegeneration and disease, to understand if and how structurally different fibrils might cause different synucleinopathies (Peelaerts et al., 2015). Recently, several groups have implemented amplification methods inspired from the prion protein field allowing the generation of fibrillar assemblies templated by pathogenic aSyn (Fenyi et al., 2019; Lövestam et al., 2021; Shahnawaz et al., 2020; Strohäker et al., 2019; Van der Perren et al., 2020). For this, two different protocols have been used, termed Protein Misfolding Cyclic Amplification (PMCA, (Saborio et al., 2001)) and real-time quaking-induced conversion (RT-QuIC, (Atarashi et al., 2011; Wilham et al., 2010)). These two fibril amplification methods differ significantly. In PMCA, fibrils are fragmented by sonication, and fibril fragments are subsequently elongated in the presence of monomeric aSyn and in the absence of any added ligands. In contrast, in RT-QuIC, fibrils are fragmented by shaking, which is performed in the presence of the ligand Thioflavin T (ThT). ThT thereby shows fluorescence upon binding to fibrils, so that the fluorescent readout of the signal from ThT can be used to follow the fibrillization process in real time. However, any ligands that bind to and thereby stabilize fibrils, risk to displace a reaction equilibrium towards the fibrillar structures that show the highest affinity for the ligand. Indeed, it has been shown that ThT impacts and accelerates the process of fibril formation and promotes more toxic oligomeric and filamentous structures, and that it influences the general structural arrangement of aSyn (Kumar et al., 2017). Lövestam et al. recently reported RT-QuIC-derived aSyn fibrillar assemblies that possessed alternate structures to the purified sample they originated from (Lövestam et al., 2021).

Here, we report the structure of aSyn fibrillar assemblies that were obtained by PMCA, using as seeds three PD and two MSA whole brain homogenates. PMCA was carefully adjusted to conditions, under which no seeding/aggregation of monomeric aSyn was observed, when brain homogenates of three control patients were used. Our determined aSyn fibril structures are novel and are denoted Polymorphs 6a, 6b and 6c. These polymorphs differ from all previously reported aSyn fibril structures, but show partial resemblance to the structure of the P-Y39 aSyn fibrillar polymorph (PDB ID 6L1T;(Zhao et al., 2020)).

Our new aSyn polymorphs have a folded core in a greek-key-like beta-sheet motif (residues 1-100), while the C-terminal flexible region (residues 124-135) is facing outwards. The reported structures differ from each other, but all share a «hook»-like hairpin motif around the NAC region, what we refer to as dragon-head shape. Our PMCA generated aSyn structures from PD and MSA patients contribute to a better understanding of the structural basis of different synucleinopathies.

## Results

### Amplification of aSyn fibrils from PD and MSA patient brain

Brain tissues obtained at autopsy from patient tissue donors who had succumbed to PD or MSA (**Table 1**) were used to amplify *E.coli* expressed and purified aSyn protein by PMCA (**Figure 1**, **Figure 1—figure supplement 2**). Briefly, 2% brain homogenates were diluted in the presence of 100μM recombinant monomeric aSyn in identical buffer conditions. Four successive amplification cycles were performed where each reaction was diluted 20 (MSA) or 100 (PD) fold in fresh 100μM recombinant monomeric aSyn. PMCA amplification was performed using post-translationally modified brain aSyn (possible phosphorylation on residues 39, 59, 64, 72, 81, 129; ubiquitination or lysine acetylation) to seed fibril growth of non-phosphorylated recombinant aSyn protein. The percentage of these modifications in sarkosyl-extracted filaments from human brain has been reported to be of low abundance for most residues except K80 ubiquitination (Schweighauser et al., 2020). These post-translational modifications may influence the PMCA-resulting fibrillar species.

**Table 1.** Patients clinical and histopathological details. 1) (Van der Perren et al., 2020) ; 2) No.: Case number of the patient; 3) PDD: Parkinson’s disease with dementia; 4) LBD: Lewy body disease; 5) DBS: Deep Brain Stimulation

| Case n° reported in ref<sup>1</sup> | No.<sup>2</sup> | Sex | Age at onset | Duration (years) | Genetics | Cause of death | Clinical diagnosis | Neuropathological diagnosis |
| --- | --- | --- | --- | --- | --- | --- | --- | --- |
| PD3 | PD405 | M | 67 | 15 | / | Not reported | Idiopathic PDD<sup>3</sup> | LBD<sup>4</sup>-Neocortical (aSyn, Braak VI); Tau Braak II, Thal phase ≥ 3 |
| PD1 | PD258 | M | 52 | 17 | PARK2 | Not reported | Tremulous parkinsonism with progression to PDD | LBD-Neocortical (aSyn, Braak VI); Tau Braak I, Thal phase 2 |
| PD4 | PD523 | F | 68 | 18 | / | Old age | PD | LBD-Neocortical (aSyn, Braak VI); Tau Braak I, Thal phase ≥ 3 |
| MSA6 | PD363 | F | 61 | 13 | / | Aspiration pneumonia | MSA bilateral pallidal DBS<sup>5</sup> (with poor response) | MSA; Tau Braak I, Thal phase 1 |
| MSA5 | PD333 | M | 52 | 8 | LRRK2 | Respiratory failure, pneumonia | Parkinsonism MSA questioned | MSA; Tau Braak I, Thal phase ≥ 3 |

**Figure 1.**
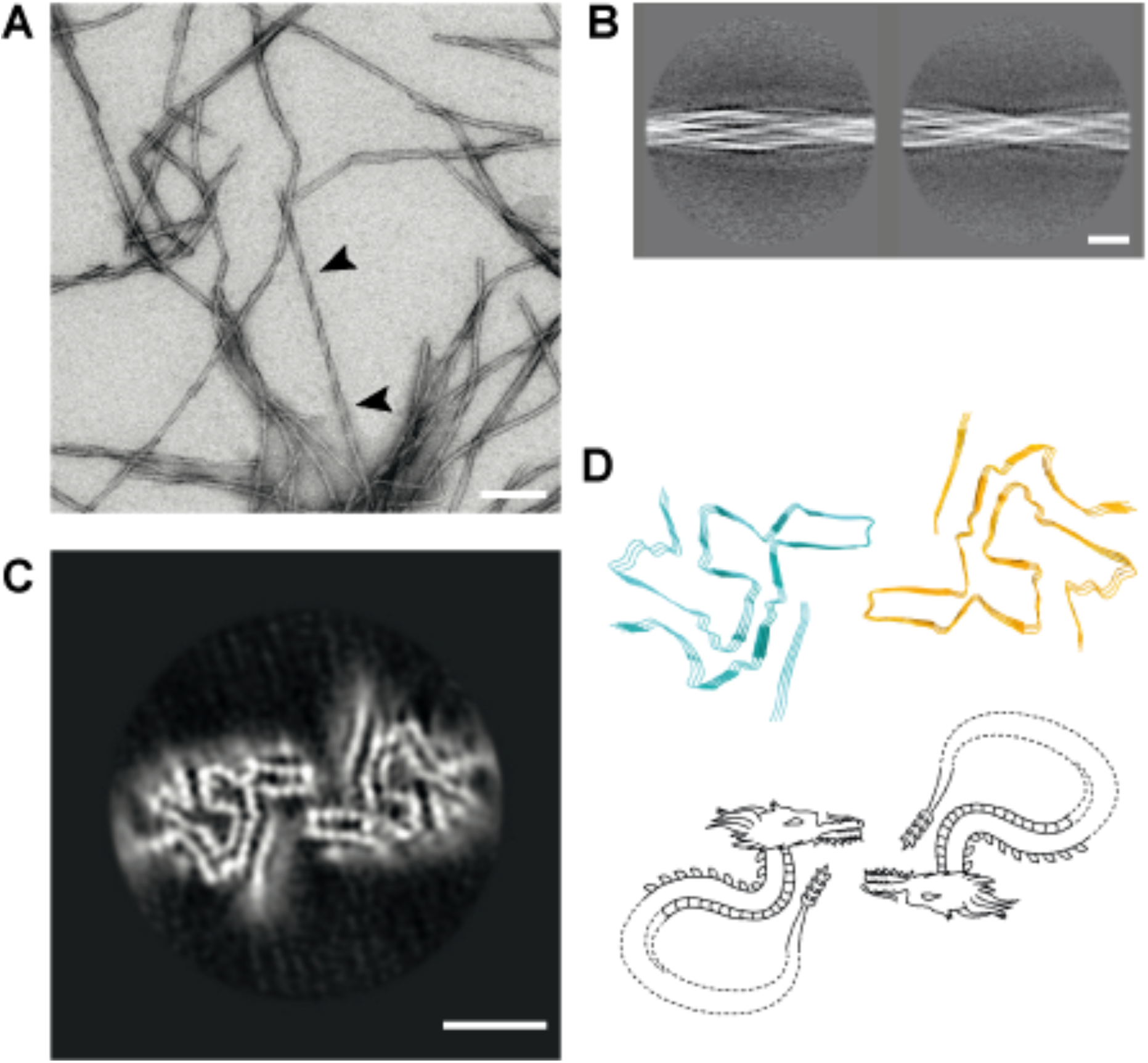
General overview of aSyn fibrils seeded by patient brain homogenate. (**A**) Negative stain (uranyl acetate) TEM micrograph depicting fibrils formed by the PMCA process using PD patient brain homogenate. Clearly visible paired helical filaments are indicated by arrowheads. The presence of non-helical fibrils is artifactual and caused by uranyl acetate staining. All fibrils appear helical in the cryo-EM. (**B**) Three times binned two-dimensional particle averages indicating a length for a 180° twist (crossover) of approximately 800Å. Scale bar = 100 Å. (**C**) The cross-section of one of the determined novel folds seeded from a PD patient brain. Another novel fold for PD and one for MSA are shown in Figure 1 Suppl. 1. Scale bar = 50 Å (**D**) Traced backbone allows for early fold estimation with a new overall structure, resembling a dragon. For Polymorph 6a, a head-to-head interface (tongue-to-tongue) between individual dragon-shaped protofilaments is visible. The other Polymorph 6b and 6c instead show a head-to-ear arrangement, with opposite faces.

As a control, brain homogenates from patients who had died from other causes and without neurodegenerative lesions were used, and no fibril formation was observed after PMCA, as judged by electron microscopy, sedimentation assay and ThT fluorescence.

As a further control, PMCA was applied under the same conditions as above but in the absence of brain extracts, yielding no fibril formation. However, when PMCA in the absence of brain extracts was carried out for extended times (beyond 5 hours), fibrils of the Polymorph 2a/2b type (Guerrero-Ferreira et al., 2019) were obtained.

The PMCA amplified samples from PD and MSA patients show a repeated helical motif of 110nm +/− 15nm, a width of 22+/−2nm, a width at crossover of 13+/−2nm and are over 1 μm long (**Figure 1 panel A**). As control, well characterized *in vitro* generated aSyn fibril seeds (Polymorph 2a/2b fibrils and aSyn Ribbons, (Bousset et al., 2013)) were amplified with PMCA to demonstrate amplification independence from the PMCA reaction buffer composition (**Figure 1–figure supplement 2**). The fibrils obtained from Polymorph 2a/2b seeds display a stiff appearance, with a central groove every 120 nm characteristic of Polymorph 2a/2b morphology. The ribbon amplified filaments produce ribbon filaments, devoid of repeated helical signature with a central groove running along the filament axis, typical to ribbons (**Figure 1—figure supplement 2**). Limited proteolytic patterns (i.e fingerprints) are identical before and after PMCA for Polymorph 2a/a and ribbons (**Figure 1—figure supplement 2**). Altogether, these results indicate that PMCA as used here can produce fibrils with characteristics related to the seeds.

### aSyn fibrils amplified from PD patient brain homogenates display two filament structures, with different protofilament interfaces

aSyn fibrils amplified from PD patient brain homogenates were first imaged by transmission electron microscopy (TEM) after negative staining with uranyl acetate. A pronounced helical motif characterized those fibrils (**Figure 1 A**). The fibrils were then analyzed by cryo-electron microscopy (cryo-EM) and optimized for high-resolution imaging. For this, samples were admitted to glow-discharged fenestrated carbon grids, plunge-frozen and transferred into a 300kV Titan Krios cryo-EM instrument. Images were recorded as dose-fractionated movies and analyzed by helical image processing routines, using the RELION software. Image processing of PD amplified samples revealed two types of filaments with equal abundance of 22’751 and 24’116 individual particle segments after 3D classification. Image acquisition and image processing data are summarized in **Table 2**. Two 3D maps at 3.97Å and 4.37Å resolution were obtained, and their respective filament structures named Polymorph 6a and 6b. The two polymorphs have a common kernel and differ in their protofilament packing. Polymorph 6a has a two-fold symmetry, while the Polymorphs 6b is asymmetric (**Figure 2—figure supplement 1**, **Figure 3**).

**Table 2.** Cryo-EM structure determination and model statistics

| Dataset | PD Polymorph 6a | PD Polymorph 6b | MSA Polymorph 6c |
| --- | --- | --- | --- |
| **Data Collection** |  |  |  |
| Nominal magnification | 165'000 | 165'000 | 165'000 |
| Pixel size (Å) | 0.82 | 0.82 | 0.82 |
| Defocus range (μm) | 0.8 - 2.1 | 0.8 - 2.1 | 0.8 - 2.1 |
| Voltage (kV) | 300 | 300 | 300 |
| Exposure time (s per frame) | 0.2 | 0.2 | 0.2 |
| Number of frames | 40 | 40 | 40 |
| Total dose (e/Å<sup>2</sup>) | 68 | 68 | 68 |
| **Reconstruction** |  |  |  |
| Box size (pixels) | 300 | 384 | 384 |
| Inter-box distance (pixels) | 35 | 35 | 38 |
| Micrographs | 1256 | 1256 | 3788 |
| Automatically picked fibrils | 4517 | 4517 | 25255 |
| Initial extracted segments | 280740 | 280740 | 616467 |
| Segments after 2D classification | 148051 | 148051 | 614531 |
| Segments after 3D classification | 22751 | 24116 | 255265 |
| Resolution after 3D refinement (Å) | 4.17 | 5.25 | 2.96 |
| Resolution after post processing (Å) | 3.97 | 4.37 | 2.67 |
| Estimated map sharpening *B*-factor (Å<sup>2</sup>) | -100 | -131.96 | -60.87 |
| Helical rise (Å) | 4.83 | 4.67 | 4.85 |
| Helical twist (°) | -0.81 | -0.9 | -0.99 |
| **Atomic model** |  |  |  |
| EMDB code | EMD-13023 | EMD-13024 | EMD-13027 |
| PDB code | 7OQ0 | 7OQ3 | 7OQ5 |
| Model resolution (Å)<br>FSC threshold | 4.2<br>(0.143) | 4.0<br>(0.143) | 2.65<br>(0.143) |
| Model resolution range (Å) | 4.4 | 4.0 | 2.70 |
| Map sharpening *B*-factor (Å<sup>2</sup>) | 95 | 120 | 30 |
| Model composition |  |  |  |
| Non-hydrogen atoms | 4374 | 4530 | 4329 |
| Protein residues | 648 | 669 | 639 |
| Ligands | 0 | 0 | 0 |
| *B*-factors (Å<sup>2</sup>) (non-hydrogen atoms) |  |  |  |
| Protein | 148 | 122 | 34.7 |
| Ligand | - | - | - |
| R.m.s. deviations |  |  |  |
| Bond lengths (Å) | 0.029 | 0.011 | 0.013 |
| Bond angles (°) | 3.62 | 2.5 | 2.79 |
| Validation |  |  |  |
| MolProbity score | 2.17 | 2.1 | 2.20 |
| Clashscore | 11.54 | 10.8 | 13.1 |
| Poor rotamers (%) | 0.0 | 0.0 | 0.98 |
| Ramachandran plot |  |  |  |
| Favored (%) | 88.5 | 88.7 | 89.3 |
| Allowed (%) | 9.6 | 10.4 | 9.8 |
| Disallowed (%) | 1.9 | 0.9 | 0.9 |

The density map reveals a central hairpin motif protruding from circularly concentric beta strands (**Figure 2 B**). The hairpin motif resembles a previously reported motif within *in vitro* generated fibrils obtained from Y39 phosphorylated aSyn protein (PDB 6L1T, residues 52-66). The latter was therefore used to initiate a manual building of aSyn structures inside the map. The root mean square deviation (RMSD) between the final model and 6L1T hairpin calculated on CA atoms coordinates for residues 52 to 66 is less than 1Å.

**Figure 2.**
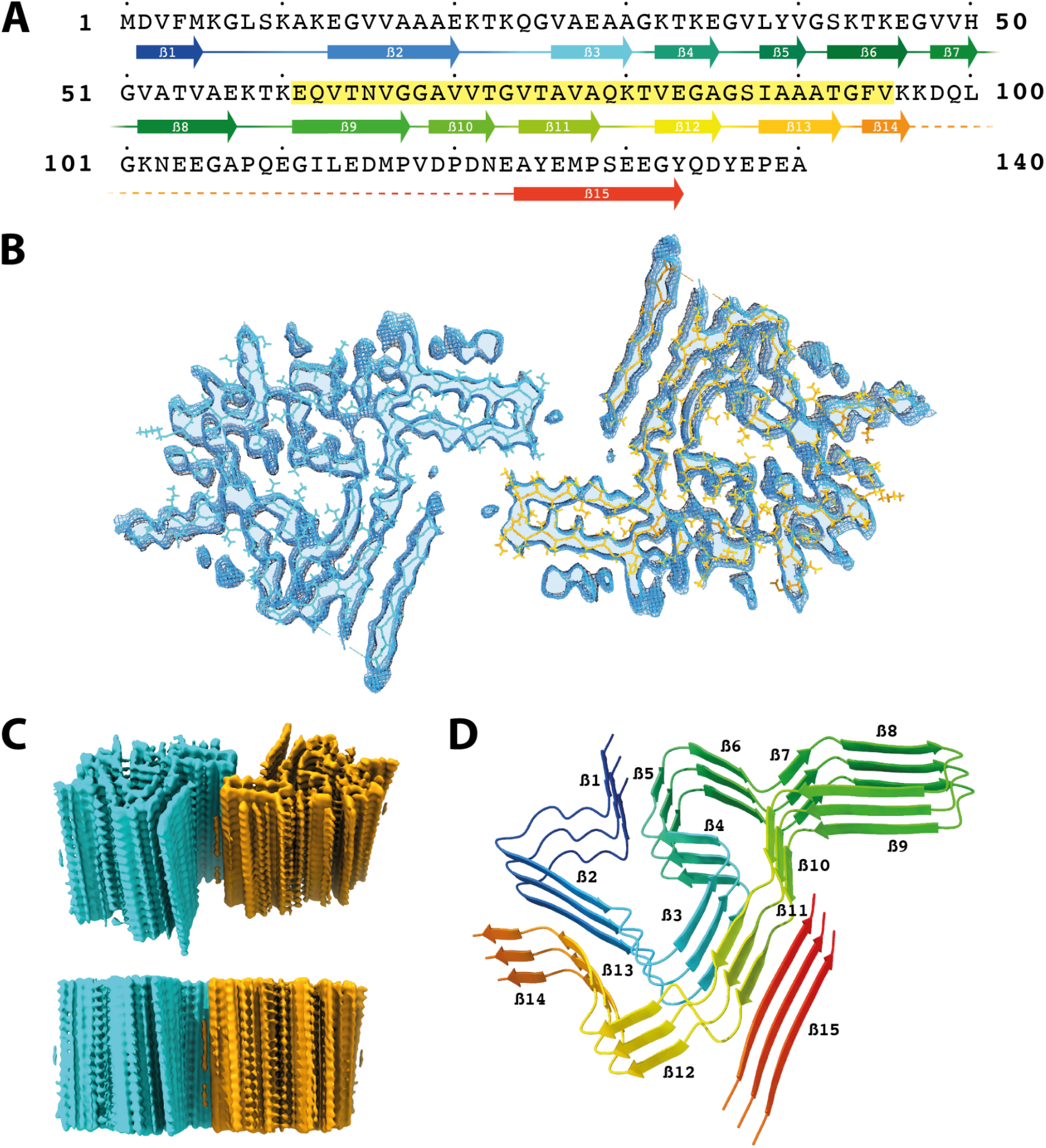
General structure of Polymorph 6a aSyn filaments seeded by PD patients brain homogenates. (**A**) Sequence of full length aSyn with the distribution of β-strands from blue (β1) to red (β15). Missing densities that do not allow establishing connectivity is marked with a dotted line (residues 98-123). The NAC-region is highlighted in yellow in the single letter amino acid residues code. (**B**) Slice through the obtained map in a central region of the reconstruction of Polymorph 6a with superimposed amino acid backbone for both protofilaments (orange and blue). A full connectivity can be observed from residue 1-97 and 124-135 resulting in a tongue-to-tongue interface of the two dragon folds. (**C**) Structural view at an angle indicates the fold interface, twist and protein stacking. The direct side view along the filament axis delivers no clear strand separation. However, typical amyloid spacing remains detectable. (**D**) β-strand distribution based on protein sequence and final model building in the density map. Residues 97-123 are not displayed as no connectivity can be observed.

Polymorph 6a comprises residues M1 to K97 and A124 to Q134, while Polymorph 6b comprises residues M1 to L100 and A124 to D135 (**Figure 2 B**, and **Figure 2—figure supplement 1 A)**. The aSyn overall fold is similar in Polymorph 6a and 6b, its cross-section is made of 15 beta-strands and resembles a dragon head (**Figure 1 D**, and **Figure 2 D)**. The central hairpin consisting of residues 52 to 66 constitutes the front part of the dragon head. Residues 32 to 51 constitute the back part of the dragon head. Residues 1 to 31 and 67 to 100 form the elongated dragon body. Its tail faces its chest and is composed of residues 124 to 135 (**Figure 2 B**, and **Figure 2—figure supplement 1 C)**. The protruding hairpin 52-66 is connected to the remainder of the polypeptide chain through a Glycine hinge composed of G47 and G67-G68, which introduces a pivot between the hairpin and the remainder of the molecule (**Figure 3—figure supplement 1 A**). The N-terminal part of aSyn (residues 20 to 47) folds into a 4 stranded S-shaped structure that creates two hydrophilic holes containing the side chains of residues K21, K23, E28 T33, E35 and residues K32, K34, Y39, E46. In Polymorph 6a, the N-terminal residues 1 to 20 compose a third hydrophilic hole made of K6, D20, E35 and G36 (**Figure 3—figure supplement 1 B**). Residues 1 to 15 totally fill this cavity in Polymorph 6b. The C-terminal part (residues 70 to 100 comprising most of the NAC domain (residues 60 to 95)) wraps around the N terminus and forms numerous hydrophobic contacts with the N-terminal part of aSyn. Hydrophobic interactions of interdigitated residues stabilize the amyloid core between β1-β5, β2-β13, β3-β11 (**Figure 3**). The region 101 to 124 is not visible in the density maps. This is indicative of a domain with higher flexibility. The C-terminal tail contacts the chest of the dragon, a beta-strand of 11 residues is clearly visible in the density, it is located 60 Å apart from residues around L100. To account for the distance, this segment can only be attributed to the region containing residues 118 to 140. We tentatively assign the residues in the poor density of Polymorph 2b as we observed that this strand is flanked by two side chain residue density bulges, a breakage in the backbone compatible with a glycine residue, and a particular density shape that could be attributed to P128 (**Figure 2—figure supplement 1 C**). The two bulges could be attributed to Y125 and Y133. Altogether, we attributed this stretch to residues A124 to D135. The intersheet distance between beta-strand T72-Q79 and E126-Q134 is notably larger than any other inter beta-sheet distances (CA-CA distance, ~9Å as opposed to ~6Å for inter beta-sheet distance between beta-strand T72-Q79 and V26-G31), compatible with the presence of bulky side chains or loose interactions at the interface. S129 is solvent exposed and may therefore be post-translationally modified, e.g., phosphorylated.

**Figure 3.**
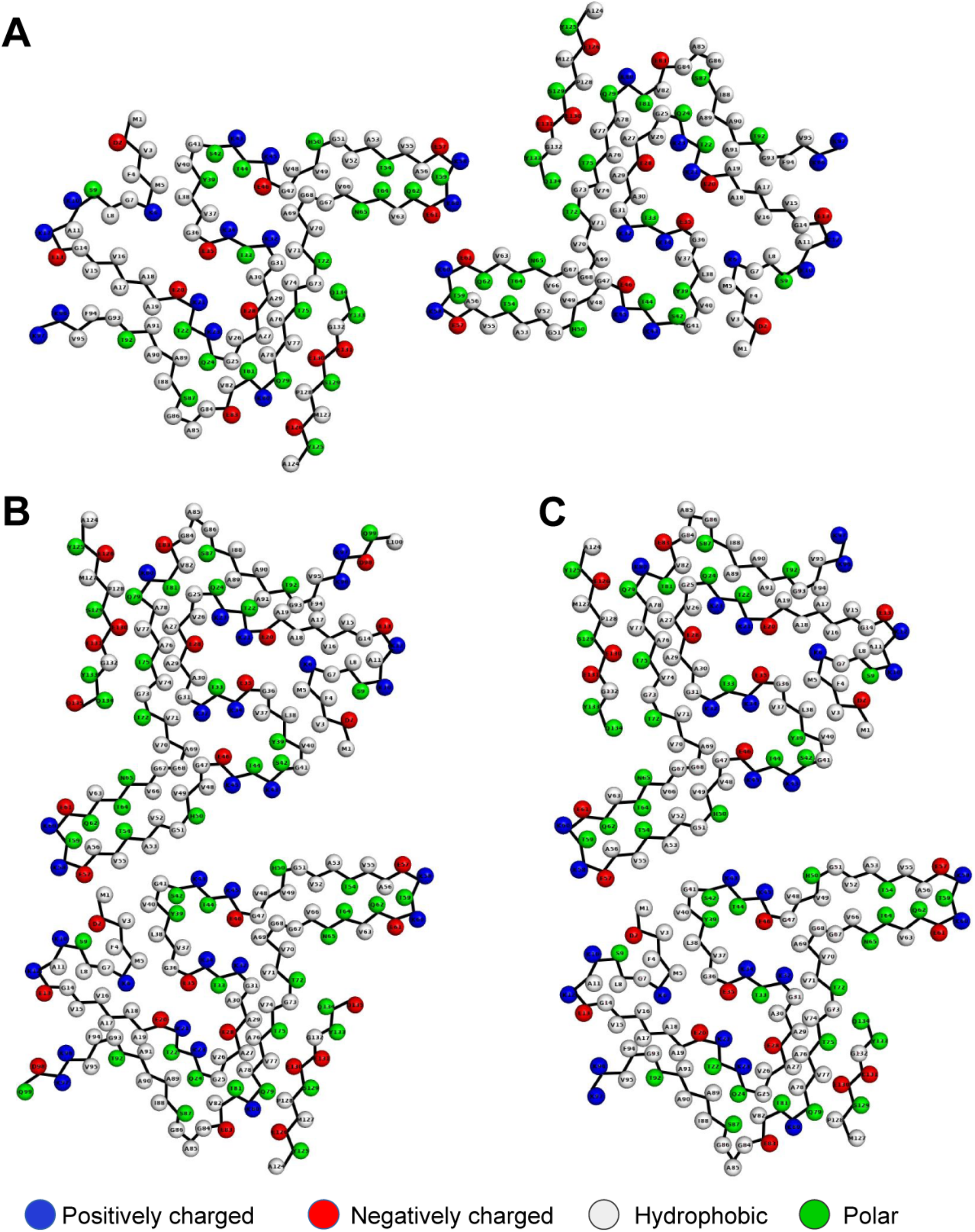
General backbone of all three fibril folds with orientation of one protofilament kept constant. Amino acid side chains are represented as spheres, colored in blue for positively charged, in red for negatively charged, in green for polar, and in white for hydrophobic residues. (**A**) First fold of PD-derived aSyn fibrils Polymorph 6a. (**B**) Second fold of PD-derived aSyn fibrils Polymorph 6b. Polymorphs 6a and b have a similar protofilament fold with a drastically different interface. Main interaction shifted towards beta-sheet 8 with interaction to beta-sheet 1 and 5 (reference figure 2). (**C**) MSA-derived aSyn fibrils fold (Polymorph 6c) display similar protofilament fold and the same interface as PD Polymorph 6b.

The interface between the two protofilaments constituting Polymorph 6a is composed of a pair of salt bridges established between K60-E61 and E61-K60 from each protofilament (**Figure 3—figure supplement 2 A**). The inter-protofilaments interface in the asymmetrical Polymorph 6b is composed of an electrostatic interaction between M1 amino terminal group and E57 side chain and Van-der-Waals interactions between residues A56-V55 and M1 side chain (**Figure 3—figure supplement 2 B**).

The structures of the amino acid stretch spanning residues 1 to 12 may account for the differences in the inter-protofilament interfaces between Polymorph 6a and Polymorph 6b. The contribution of these residues to the inter-protofilaments interface in Polymorph 6b has also an impact on the size of the surrounding hydrophilic channel (**Figure 3—figure supplement 1B**, and **Figure 3—figure supplement 2B**)

### aSyn fibrils amplified from MSA patient brain homogenates exhibit a unique structure, resembling Polymorph 6b

aSyn fibrils amplified from MSA patient brain homogenates were imaged and analyzed the same way as described for those amplified from PD patient brain homogenates. The image analysis using the RELION software revealed the presence of a single type of filaments. A 3D map at 2.67 Å was obtained and the structure was named Polymorph 6c (**Table 2**). The structure of Polymorph 6c resembles that of Polymorph 6b, and displays an asymmetric protofilament organization (**Figure 2—figure supplement 1B, Figure 3C** and **Figure 3—figure supplement 2 C**).

Polymorph 6c comprises residues M1 to K97 and M127 to Q134 in one protofilament and residues M1 to K97 and residues A124 to Q134 in the second. Polymorph 6c differs from Polymorph 6b by a 4Å larger protofilament interdistance. The monomeric aSyn structure in Polymorph 6c is nonetheless highly similar to the protofilaments in Polymorph 6b with an RMSD over all determined CA atoms of 1.5 Å.

### Comparison of Polymorph 6, de novo assembled Phospho-Y39 aSyn fibrils, de novo assembled aSyn fibrils and brain extracted MSA filament folds

aSyn filamentous polymorphs with asymmetric protofilament organization obtained upon amplification from PD (Polymorph 6b) and MSA (Polymorph 6c) patient brain homogenates exhibit very limited structural differences. The aSyn molecule structures of the two protofilaments within each polymorph differ by an RMSD over 110 CA atoms of 2.7 Å for Polymorph 6b, and 2.9 Å for Polymorph 6c (**Figure 4 B, C**). A slight rotation of the protruding hairpin relative to the compact core around the glycine hinge distinguishes Polymorphs 6b from 6c. Significant differences are observed between the symmetrical (6a) and asymmetrical (6b and 6c) protofilament organization. aSyn’s 15 first amino acid residue stretch appears more compactly folded in Polymorph 6b and 6c (**Figure 4 B and C)**, as compared to Polymorph 6a (**Figure 4 A)**. The maximum CA displacement is 6Å and the overall deviation between the symmetrical and asymmetrical polymorphs remains limited below 2.9 Å (**Figure 4D**). Surprisingly, Polymorphs 6b and 6c only differ by a slight increase of the inter-protofilament distances (**Figure 4 E**).

**Figure 4.**
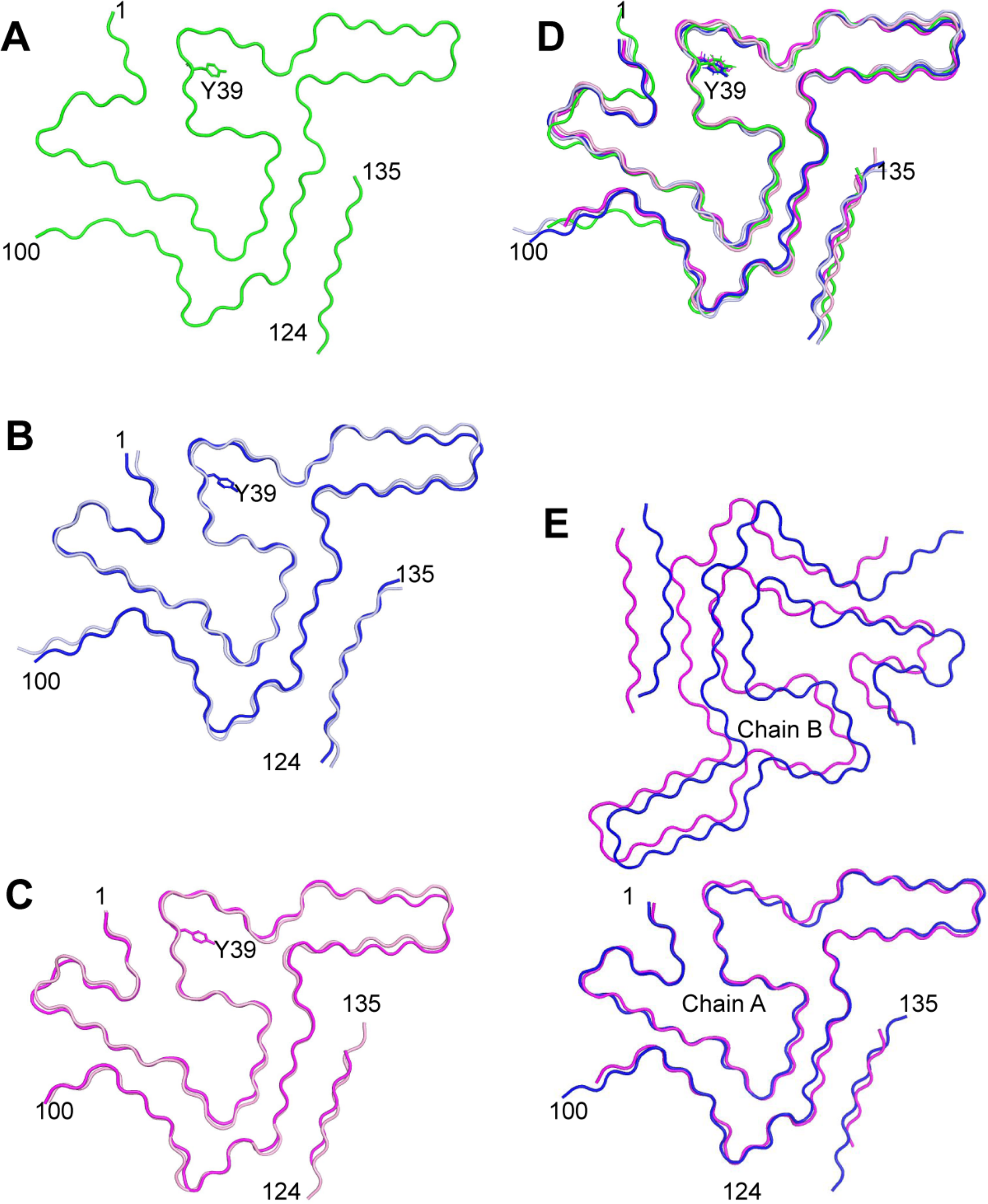
Polymorph 6 structure comparisons. (**A**) Backbone structure of Polymorph 6a (green). (**B**) Superimposition of Polymorph 6b chain A (protofilament 1, blue) and B (protofilament 2, grey), RMSD value of 2.7 Å over 110 CA. (**C**) Superimposition of Polymorph 6c chain A (protofilament 1, purple) and B (protofilament 2, pink). RMSD value of 2.9 Å over 110 CA. (**D**) Superimposition of Polymorph 6a (green), 6b (chain A, blue and B, grey), and 6c (chain A, purple and B, pink). (**E**) Superimposition of Polymorph 6b (in blue) to Polymorph 6c (in purple), upon aligning the A chains (protofilaments 1).

The structure of Phospho-Y39 aSyn fibrils (PDB# 6L1T) assembled de novo, shows 4 lysine side chains (K21, K32, K34 and K43) surrounding the strongly negatively charged phosphate group on Y39 hydroxyl group. This results in a conformational rearrangement yielding a hydrophilic channel, running along the protofilament core, that appears much wider than any of the 2 channels resulting from Y39 hydroxyl group interaction with K34 side chain in Polymorphs 6a, 6b and 6c (**Figure 3 – figure supplement 1B**). The stretch of amino acid residues 52-66 appears to adopt highly conserved structures in Polymorphs 6a, 6b, 6c, de novo assembled phospho-Y39 aSyn fibrils and de novo assembled human full-length aSyn (PDB 6ssx), despite being in a different tertiary environment in the latter case, with RMSDs below 0.59 Å between all polymorph 6 structures and PDB # 6l1t and RMSDs below 1.5 Å, between all polymorph 6 structures and PDB # 6ssx (**Figure 4—figure supplement 1 A, B, D**). The structure of the stretch formed by amino acid residues 70-97 appears also conserved in Polymorphs 6a, 6b, 6c and de novo assembled Phospho-Y39 aSyn fibrils, with RMSDs below of 3.0 Å (**Figure 4—figure supplement 1 C**). In contrast with what we report for Polymorphs 6a, 6b and 6c, the aSyn C-terminal polypeptide stretch spanning residues 124 to 134 appears unresolved both in phospho-Y39 aSyn fibrils and de novo assembled human full-length aSyn.

The structures for patient brain extracted MSA filaments (PDB 6xyo) reported by Schweighauser et al. (Schweighauser et al., 2020) shared no similarity with any of the aSyn polymorph structures reported here. In 6xyo, two hairpins are observed, formed by residues 14 to 31 in one of the protofilaments and 25-46 in the other. Residues 47-99 appear to form two head-to-tail L-shaped structures, and the aSyn N- and C-terminal amino acid stretches formed by residues 1-14 and 100-140 in one protofilament, and by 1-21 and 100-140 in the other protofilament, are unresolved. This contrasts with all Polymorph 6 variants reported here, where aSyn residues 1-12 adopt a V-shaped structure, residues 98-123 are unresolved and residues 124-134 are stacked to the dragon-shape belly (**Figure 4—figure supplement 1 E**).

## Discussion

aSyn is capable of forming several structurally distinct polymorphs *de novo* (Bousset et al., 2013; Guerrero-Ferreira et al., 2020, 2019, 2018; Y. Li et al., 2018; Verasdonck et al., 2016). For this, the experimental conditions, mutations, truncations and post-translational modifications within the protein play a key role in the structural diversity of the resulting aSyn amyloids (Crowther et al., 1998; Zhao et al., 2020). It has also been shown that the intrinsic structural characteristics of distinct aSyn fibrillar polymorphs are faithfully reproduced upon elongation of preformed polymorphs in the presence of monomeric aSyn (Bousset et al., 2013). When structurally distinct pure fibrillar aSyn polymorphs were injected into the central nervous system (CNS) of model animals, distinct synucleinopathies were observed (Peelaerts et al., 2015). This suggests that distinct aSyn fibril strains with specific structure-phenotypic traits may be correlated with, or in part even causatively responsible for specific synucleinopathies. To further support this view, we exploited the capacity of pathogenic aSyn aggregates present in the brain of PD, MSA and DLB patients to recruit monomeric aSyn through PMCA, a method inspired from the amplification of pathogenic prion protein particles in brain homogenates (Saborio et al., 2001). The resulting fibrillar assemblies were shown to exhibit different shapes and characteristic proteolytic patterns indicative of specific structures (Van der Perren et al., 2020). They further triggered defined synucleinopathies upon inoculation in rodent brains and seeded to different extent the aggregation of aSyn in human dopaminergic neurons derived from induced pluripotent stem cells (iPSC) (Tanudjojo et al., 2021). This further supports a structure-phenotype relationship.

We here report the structures of three fibrillar polymorphs obtained upon repeated cycles of seeding, elongation and fragmentation of recombinant, monomeric, non-phosphorylated aSyn by phosphorylated, pathogenic aSyn aggregates present in brain homogenates of PD and MSA patients in the absence of ThT or detergents. The presented polymorphs obtained by PMCA display three novel folds, and we have shown that our PMCA method is able to maintain the structure of *de novo* formed distinct aSyn polymorphs. The folds of aSyn reported here roughly resemble that of Phospho-Y39 aSyn fibrils (PDB# 6L1T) and in part bear resemblance to that of unmodified Polymorph 2a/b aSyn (PDB# 6SSX). The shared structure spans 14 residues (residues 52-66) with Polymorph 2 and 30 additional C-terminal residues for Y39 phosphorylated aSyn (residues 52-66 and 70 to 100). Among the numerous polymorphs of aSyn structures solved, slight variations have been observed between closely related folds, indicating possible local fold remodeling. PMCA uses raw homogenized patient’s brain extract, the availability of the templates and their ability to incorporate recombinant unmodified aSyn directly determine the nature of the resulting filaments.

The fold appears conserved within the three fibrillar polymorphs we present, however the protofibrils constituting each polymorph appear organized differently. Polymorph 6a exhibits a two-fold symmetry while the two others (6b and 6c) are asymmetrical. PD brain homogenates led to the formation of equal proportions of two polymorphs (6a and 6b), while only one polymorph (6c) was obtained from MSA brain homogenates. These differences, together with the observation that amplification reactions performed in brain homogenates of age matched non-demented patients yielded no fibrils suggests that the fibrillar assemblies we generated result from a seeding process specific to PD or MSA.

Previously described patient brain extracted MSA filaments ((Schweighauser et al., 2020), PDB 6xyo, 6xyp and 6xyq) shared no fold similarity with any of the aSyn Polymorph 6 structures we report here.

Two hairpins are observed in those MSA extracted filaments, formed by residues 14–31 in one and 25-46 in the other protofilament. Residues 47-99 appear to form two head-to-tail L-shaped structures, and aSyn N- and C-terminal amino acid stretches, residues 1-14 and 100-140 in one protofilament, and 1-21 and 100-140 in the other, remain unresolved. This contrasts with our here reported polymorph 6 structures, in which aSyn residues 1-12 adopt a V-shaped structure, residues 98-123 are unresolved and residues 124-134 are stacked to the dragon-shaped belly (**Figure 4—figure supplement 1 E**).

Despite the structural differences between fibrils derived from PD and MSA patients brain homogenates, the finding that our amplified fibrils share no fold similarity with MSA filaments extracted in the presence of detergents from MSA patient brains question the faithfulness of templating. To address this and to demonstrate that PMCA under our experimental conditions is able to faithfully reproduce a given aSyn fold, we amplified two distinct *de novo* assembled aSyn fibrillar polymorphs. The resulting amplified fibrils exhibited the characteristics of the input polymorphs as judged from TEM analysis and limited proteolysis profiling (**Figure 1 – Figure supplement 2**). Amplification is performed under physiological experimental conditions in our case, in the absence of detergents commonly used to purify pathogenic aSyn aggregates from patient brains, and in the absence of fibril-binding ligands, such as ThT, that exhibit differential affinity for distinct polymorphs, thus orienting aggregation towards the species it binds most.

The structures we report are distinct from those derived from aSyn seeds extracted through centrifugation cycles and in the presence of detergents from MSA patients brain homogenates using another amplification method. One possible reason for this difference could be the method used, in particular when traces of detergents or the ligand ThT are present in the reaction mix. Another reason may come from changes in the structure of the seeds during the extraction procedure. Finally, the discrepancy could be caused by the amplification process, which may select the best seeder, if an ensemble of different templates were present among the seeds. This might have affected the attempts to amplify aSyn filaments by RT-QuIC from raw brain or semi-purified seeds (Lövestam et al., 2021). Growing evidence shows that in several neurodegeneration cases, a mix of multiple polymorphs populates diseased brain tissues (Shi et al., 2021). Further characterization of these filaments will provide insight into their respective role in pathology. In any case, hope prevails that at least one of these several structures might turn out to be the key to understanding the mechanistic processes within neuronal cells that lead to neurodegeneration, so that pharmaceutical strategies to modify the synucleinopathies can be developed.

## Conclusions

We present the ultrastructure of three aSyn fibril polymorphs, which were PMCA amplified from human brain tissue from Parkinson’s disease (Polymorphs 6a and 6b) and Multiple System Atrophy (Polymorph 6c) patient tissue donors. The employed PMCA amplification protocol did not involve any usage of sarkosyl or other detergents, fibril-binding ligands, precipitation or drying at any step. PMCA seeded from control brain tissue did not result in fibril formation, and PMCA-derived fibrils from structurally distinct *in vitro* generated fibrils resulted in fibrils of different structures. All three here reported fibril polymorphs for PD and MSA are composed of two protofibrils that bear a partial resemblance to the *in vitro* fibrils formed by Y39-phosphorylated aSyn protein. Further work will be needed to reliably correlate fibril strain with disease subtypes, so that a putatively causative role of specific fibril polymorphs for different synucleinopathies can be identified.

## Materials and Methods

### Human brain

Frozen brain tissues from patients who succumbed from PD and MSA, as well as from age-matched healthy control tissue donors, were obtained at autopsy from three PD and 2 MSA patients from UK Brain Bank (Imperial College London, UK). The clinicopathological description of the 5 patients is summarized in **Table 1**. The cingulate cortex was isolated from the brains of the patients with PD. The cerebellum was obtained from the brains of the patients with MSA. Both regions were also extracted from two healthy control brains.

### Brain tissue homogenization

Frozen brain tissues were weighed in falcon tubes (15 or 50 ml depending on the total weight). The samples were diluted five times in PMCA buffer (150 mM KCl, 50 mM Tris–HCl pH 7.5) to obtain a homogenate at 20% (weight:volume). The homogenization was performed by sonication using the SFX 150 Cell Disruptor sonicator with a 3.17 mm microtip probe (Branson) for 1 min, with 10 s pulses followed by 10 s pauses in a biosafety level 3 environment (BSL-3). The homogenates were aliquoted and immediately frozen in liquid nitrogen before storage at −80°C.

### Recombinant protein

Monomeric aSyn was expressed and purified as described in (Bousset et al., 2013) and (Gath et al., 2014). Briefly, recombinant, wild-type aSyn was expressed in *E. coli* strain BL21(DE3), transformed with the expression vector pET3a (Novagen) encoding wild-type, full-length aSyn. Expression was induced by 0.5 mM IPTG for 2 hr, when the bacteria grown in LB medium at 37°C had reached an optical density of 1.0 at 660 nm. Soluble, monomeric aSyn was purified from the bacterial lysate by ammonium sulfate precipitation, followed by PEI nucleic acid precipitation, and the resulting clarified lysate was loaded on DEAE sepharose resin. Fractions containing alpha synuclein were heated to 80°C for 10 minutes and spun at 10.000g. The aSyn concentration in the supernatant was determined spectrophotometrically using an extinction coefficient of 5960 M^−1^*cm^−1^ at 280 nm. Pure aSyn (0.7 mM) in 50 mM Tris-HCl, pH 7.5, 150 mM KCl was filtered through sterile 0.22 μm filters and stored at −80°C.

### Proteinase K proteolytic digestion

De novo assembled aSyn fibrils and ribbons (1.4 mg/ml) in 150 mM KCl, 50 mM Tris–HCl, pH 7.5, were treated at 37 °C by Proteinase K (3.8 μg/ml) (Roche). Aliquots were removed at 1, 5, 15, 30 and 60 minutes following addition of the protease and transferred into Eppendorf tubes maintained at 90 °C containing denaturing sample buffer (50 mM Tris–HCl, pH 6.8, 4% SDS, 2% beta-mercaptoethanol, 12% glycerol and 0.01% bromophenol blue) to arrest immediately the cleavage reaction. After incubation of each tube for 5 min at 90 °C, the samples were processed to monitor the time course of aSyn cleavage by polyacrylamide gel electrophoresis (PAGE) (15%) after staining with Coomassie blue.

### Fibril amplification by protein misfolding cyclic amplification (PMCA)

All operations were performed under biosafety level 3 (BSL-3) conditions. Brain homogenates were diluted in PMCA buffer (150 mM KCl, 50 mM Tris–HCl, pH 7.5) containing monomeric aSyn (100 μM) to a final concentration of 2% (weight:volume), equivalent to 6 mg of brain tissue, as described previously for other tissues (Van der Perren et al., 2020).The sample was split into two PCR tubes (BIOplastics, Landgraaf, The Netherlands). PMCA amplification was performed in quadruplicates for each patient using the Q700 generator and a 431MPX horn (Qsonica, Fisher scientific, Illkirch, France). The power of the horn was set to 30% of maximal amplitude. The program of amplification consisted in 15 s of sonication and 5 min pause at 31°C. Every hour, 5 μl were withdrawn from each tube and diluted in 300 μl of 10 μM of ThT. ThT was not brought into contact with the remainder of the sample. The progress of the amplification was monitored by measuring ThT fluorescence in the withdrawn samples, using a Cary Eclipse Fluorescence Spectrophotometer (Agilent, Les Ulis, France) with fixed excitation and emission wavelength at 440 nm and 480 nm respectively. Cycle 2, 3 and 4 were performed following the same protocol using 1% of the preceding cycle reaction as seeds for PD or 5% for MSA cases. The amounts of brain homogenates and PMCA amplified assemblies used in each amplification reaction were defined through an optimization study aimed at maintaining high stringency by minimizing the de novo aggregation of aSyn under our experimental conditions. The time at which an aliquot from one amplification reaction was withdrawn for a subsequent amplification reaction was also defined through an optimization study aimed at avoiding the de novo formation of aSyn fibrils.

At cycle 4, PMCA reaction products were left overnight at 37°C without shaking, to let the elongation reaction reach completion. The obtained samples were used directly for negative stain TEM or Cryo-EM imaging, without dilution or sedimentation.

### Negative staining and transmission electron microscopy

The morphology of the *de novo* assembled and PMCA-amplified aSyn assemblies was assessed by transmission electron microscopy (TEM) in a Jeol 1400 transmission electron microscope, following adsorption of the samples onto carbon-coated 200 mesh grids and negative staining with 1% uranyl acetate. Images were recorded using a RIOS camera (Gatan Inc).

### Cryo-Electron microscopy

Cryo-EM grids were prepared using a Vitrobot Mark IV (Thermo Fisher Scientific) plunge freezer from undiluted samples at room temperature and visualized immediately on an FEI Talos TEM. After determination of optimal freezing conditions, final cryo-EM grids were prepared from 3μl of undiluted sample applied onto 45s glow-discharged, 300 mesh, copper Quantifoil grids (R2/1) followed by rapid plunging into liquid ethane to freeze.

Micrographs were acquired on a Titan Krios G1 (Thermo Fisher Scientific) transmission electron microscope, operated at 300 kV and equipped with a Gatan Quantum-LS imaging energy filter (GIF, 20 eV zero loss energy windows; Gatan Inc). Images were recorded on a K2 Summit electron counting direct detection camera (Gatan Inc) in dose fractionated counting mode (40 frames, 0.2 seconds exposure time per frame) using the SerialEM software (Mastronarde, 2005) at a pixel size of 0.82 Å for the physical pixels and a total dose of ~68 electrons per square Angstrom (e-/Å2) for each micrograph. Recorded images were directly processed using FOCUS (Biyani et al., 2017) to monitor data quality during imaging. Within FOCUS, all micrographs were subjected to drift-correction and dose-weighting using MotionCorr2 (Zheng et al., 2017) and CTF estimation using Ctffind4.1 (Rohou and Grigorieff, 2015).

### Image processing

Image processing and helical reconstruction were carried out with RELION 3.1, following the methods described for amyloid structure determination (Scheres, 2020). Individual filaments were selected using crYOLO (Wagner et al., 2019) using a trained model for each of the datasets. For all reconstruction and processing steps, detailed information is given in **Table 2**. Filaments were traced using a box size of 200 pixels and an inter-box distance of 35 pixels. The box coordinates were imported into RELION and used for consecutive segment extraction, in a first iteration with 4-times binned particles to an effective size of 2.46 Å per pixel. After 2D classification with a regularization parameter of T=2, the cross-over distance of the fibrils was measured on the 2D class averages that showed a clear alignment (**Figure 1B**). Measurements resulted in an approximate distance of 800Å for an 180° twist for all fibrils. The *relion_helix_inidmod2d* function was used to generate an initial model with application of the cross-over distance measured and assuming a helical rise of 4.75Å. The resulting 3D map was used in the consequential processing steps as a low-pass filtered starting reference.

#### Fibrils seeded from PD tissue

Initially, 3D classification with four classes and the 60Å lowpass-filtered reference separated the dataset into two distinct subforms with a respective particle count ratio of approximately 50:50 (70’519:77’532 particles). For further processing, the subgroups were treated separately. Re-extraction of the specific particles was performed with an unbinned box size of 352 pixels for group 1 and 384 pixels for group 2 at an effective size of 0.82 Å per pixel. The map from the previous step was resized and rescaled for the respective group and used as a reference. Repetition of 3D classification on the unbinned particles with fixed helical parameters for twist and rise (−1° and 4.75Å), with four classes for group 1 and three classes for group 2, resulted in a map where the general backbone of the respective fold for each group became visible (**Figure 1 D**, and **Figure 1–figure supplement 1**). Following iterative steps of *3D auto-refine* and *ctf_refine* in RELION 3.1 on the best aligned class with C1 symmetry optimization of helical twist and rise resulted in two structures with overall resolutions of 4.17 Å (Polymorph 6a, **Figure 2B**) and 5.25 Å (Polymorph 6b, **Figure 2–figure supplement 1 A**). Post-processing with soft-edge masks and an estimated map sharpening B-factors of −100 Å^2^ (manually limited) and −131.96 Å^2^ (unrestricted), respectively, improved the maps to 3.97 Å and 4.37 Å.

#### Fibrils seeded from MSA tissue

The crYOLO traced particles were extracted using a 384 pixel box size. Subsequent 2D classification and selection of classes that indicated a clear strand separation resulted in 614’531 particles. For 3D classification, the rescaled and low-pass filtered (60 Å) initial model generated from the 2D binned particles was used, using a helical reconstruction with fixed symmetry for twist and rise (−1° and 4.75Å). Out of a total of three classes one clear fold similar to Polymorph 6b became evident (**Figure 1–figure supplement 1 B**). A second class average was detected with low resolution alignment. The classes were separated into two respective groups with 255’665 and 90’755 particles each, all further processing was performed separately on each class. Additional 3D classification removed noise and reduced the remaining particles to 141’067 and 80’267 particles, respectively. While for group 1 a better alignment was achieved, group 2 did not show any significant improvement in map quality. Finally, iterative steps of 3D refinement for local helical symmetry and CTF refinement as well as polishing resulted in a map of 2.67 Å after post-processing with soft-edge mask and an estimated map sharpening B-factor of −60.8Å for group 1, resulting in Polymorph 6c (**Figure 2–figure supplement 1 B**). Similar processing of group 2 did not yield any interpretable results and we thus cannot reliably document this fold in MSA.

### Model building and refinement

Models for the PD fibril Polymorph 6b and MSA fibril Polymorph 6c were built into the RELION local resolution-filtered maps with COOT (Emsley and Cowtan, 2004), based on the structure of the aSyn filament with a phosphorylated Tyrosine 39 (PDB 6L1T)(Zhao et al., 2020). The 6L1T structure was manually docked into the 3D maps using the hairpin as marker, and the remainder of the protein with a modified conformation was rebuilt manually into the densities. The backbone trace from residues 1 to 99 was built into the Polymorph 6a map, but the lack of a clear 4.7 Å Z-axis separation did not allow a precise Z axis positioning. The later determined Polymorphs 6b and 6c were iteratively used to confirm Z-axis positions in Polymorph 6a.

To build the model for Polymorph 6b, the Polymorph 6a model was docked into the 6b map and manually rebuilt. The initial model for Polymorph 6c was obtained by a rigid body fitting of Polymorph 6b protofilaments. The structures were then refined against their respective maps with PHENIX real space refinement (Afonine et al., 2018), using rotamer and Ramachandran restraints, as well as non-crystallographic symmetry and beta strand geometry constraints. Refined structures were inspected with UCSF Chimera (Pettersen et al., 2004) and ISOLDE (Croll, 2018) with in-register parallel beta strand restraints. For Polymorph 6a, C2 symmetry was applied. For Polymorph 6b and 6c, each protofilament was built independently. All structures were validated using Molprobity(Williams et al., 2018). Figures were prepared using UCSF Chimera and Pymol (The PyMOL Molecular Graphics System, Version 2.0 Schrödinger, LLC.).

## Data availability

The cryo-EM image data are available in the Electron Microscopy Public Image Archive, entry number EMPIAR-XXX. The 3D map is available in the EMDB, entry number EMD-13023, EMD-13024, EMD-13027. The atomic coordinates are available at the PDB, entry number PDB 7OQ0, 7OQ3, 7OQ5.

## Acknowledgements

We acknowledge Steve Gentelman, Imperial College, London, for providing frozen patients brain tissues. This work would not have been achieved without the support of UK Brain Bank, the patients, their families, and their caregivers that we gratefully acknowledge. We thank Kenneth N. Goldie, Mohamed Chami and Lubomir Kovacik for support in cryo-EM data collection. This work has benefited from the platform and expertise of the Electron Microscopy Facilities of I2BC, we thank Ana Andreea Arteni for screening samples with Cryo-EM. Calculations were performed at the scientific computing center of the University of Basel (sciCORE, http://scicore.unibas.ch/) and on own laboratory hardware.

## Funding Sources

*European Union’s Horizon 2020 research and innovation program and EFPIA Innovative Medicines Initiative 2 grant agreement No 116060 (IMPRiND) and the Swiss State Secretariat for Education, Research and Innovation (SERI) under contract number 17.00038*

Ronald Melki

*The EU Joint Programme on Neurodegenerative Disease Research and Agence National de la Recherche (contracts PROTEST-70, ANR-17-JPCD-0005-01 and Trans-PathND, ANR-17-JPCD-0002-02)*

Ronald Melki

*Fondation pour la Recherche Medicale (contract DEQ. 20160334896 and ALZ201912009776)*

Ronald Melki

*JiePie Award 2019*

Ronald Melki

Swiss National Science Foundation, grant CRSII5_177195

Henning Stahlberg

Domenic Burger

Swiss National Science Foundation, grant 310030_188548

Henning Stahlberg

Domenic Burger

---

**Figure 1—figure supplement 1.**
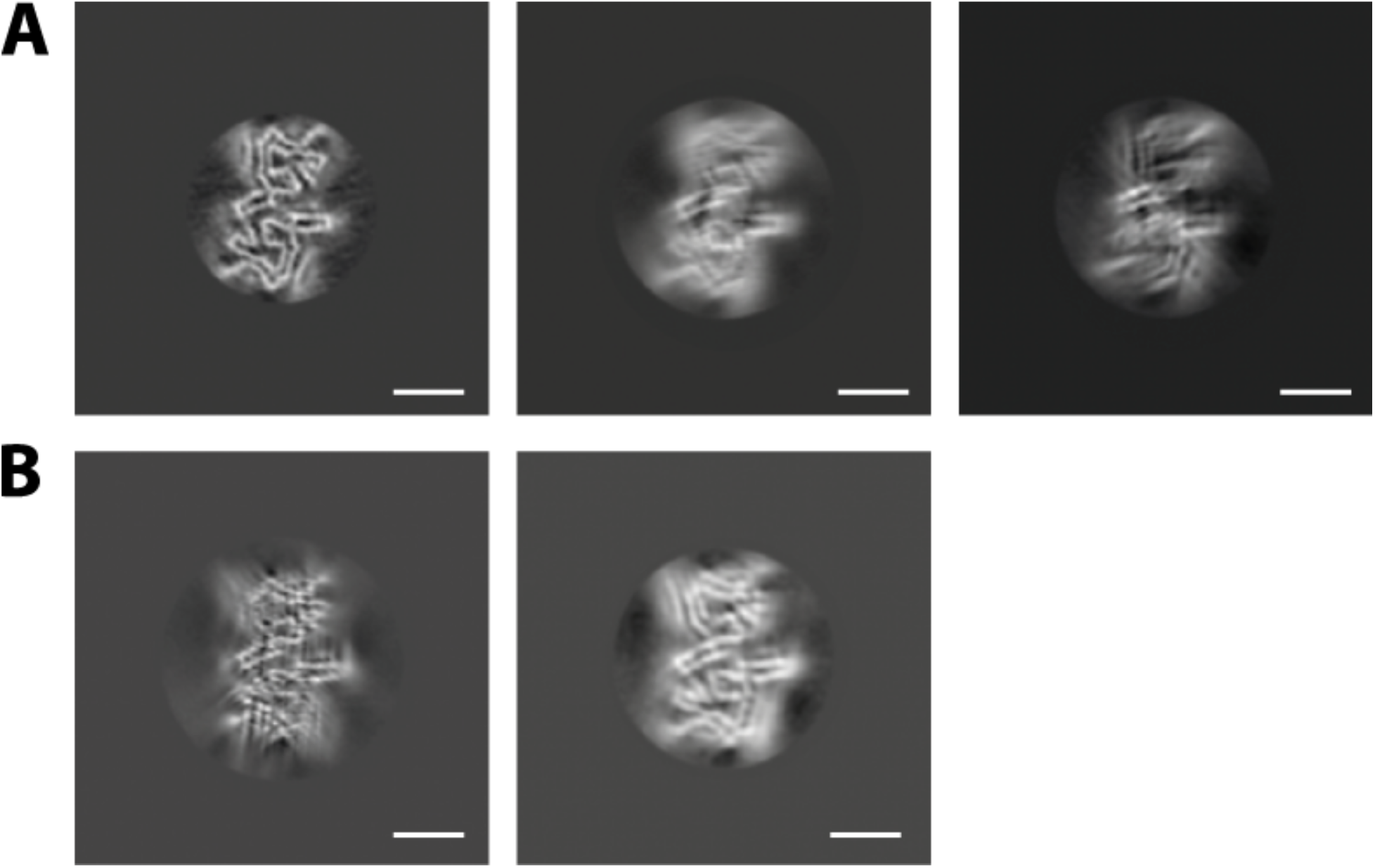
Structural overview of fibrillar aSyn folds from different PD and MSA patients. (**A**) Polymorph 6b from PD patient (PD405) brain homogenate (left), and lower-resolution maps of fibrils seeded from two additional PD patients brain homogenates (PD258 and PD523). (**B**) Polymorph 6c from two MSA patients: MSA363 (left) and MSA333 (right). For details, see Table 1. Scale bars = 50 nm.

**Figure 1—figure supplement 2.**
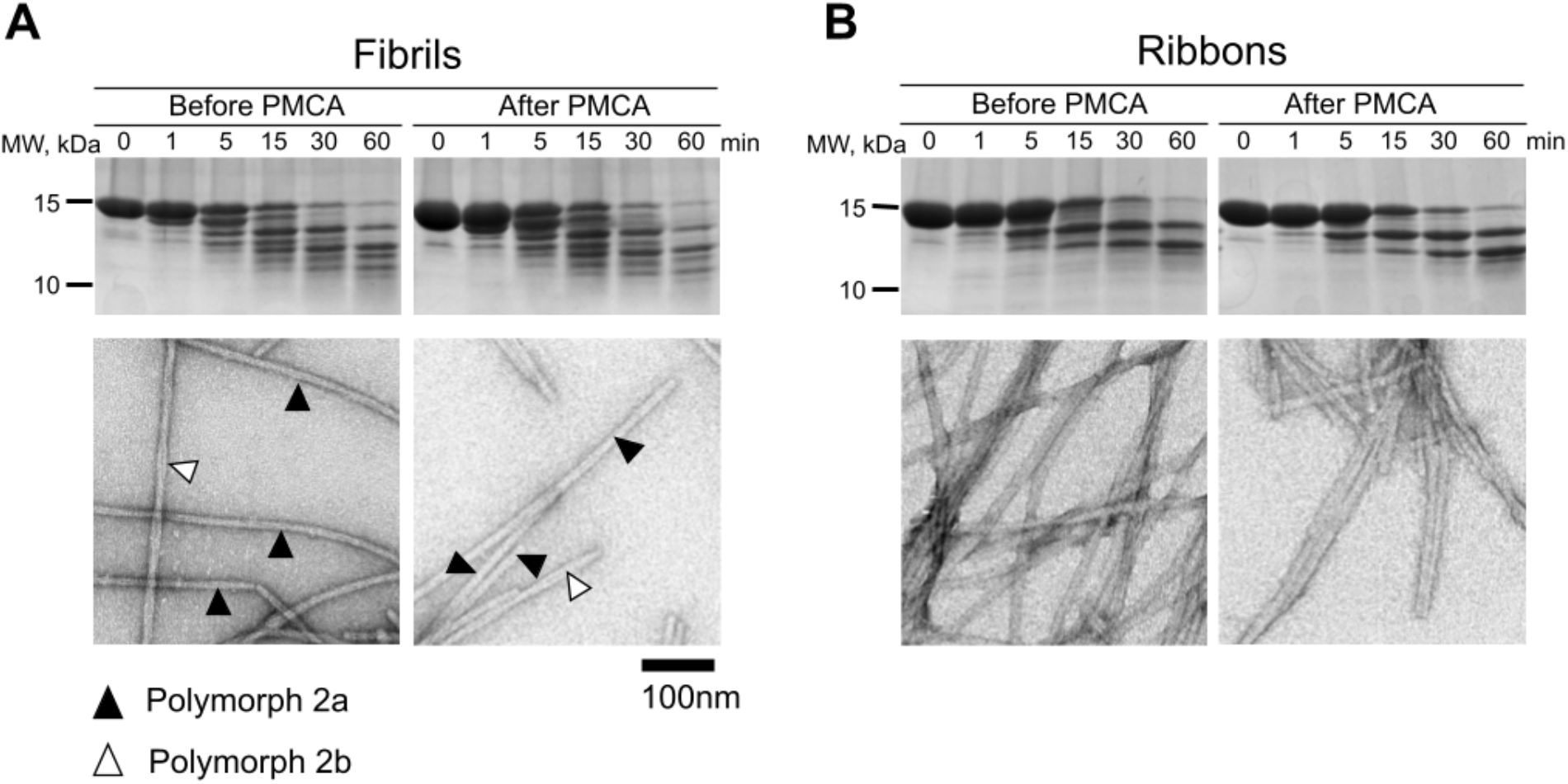
Verification of PMCA amplification fidelity, using two structurally distinct *in vitro* assembled fibrils. (**A**) “Fibrils” proteinase K proteolytic profiles before and after amplification, and negative stain transmission electron micrographs of Polymorph 2a/b filaments before and after amplification. Limited proteolysis pattern indicates a similar accessibility to protease, indicative of a common fold. Typical helical repeat signature of Polymorph 2a/b is indicated by black for Polymorph 2a, and white for Polymorph 2b arrowheads. (**B**) “Ribbons” filaments proteinase K proteolytic profiles before and after amplification, and negative stain TEM images before and after amplification. Limited proteolysis pattern indicates a similar accessibility to protease, indicative of a common fold. The typical non helical signature of ribbons is observed before and after PMCA.

**Figure 2—figure supplement 1.**
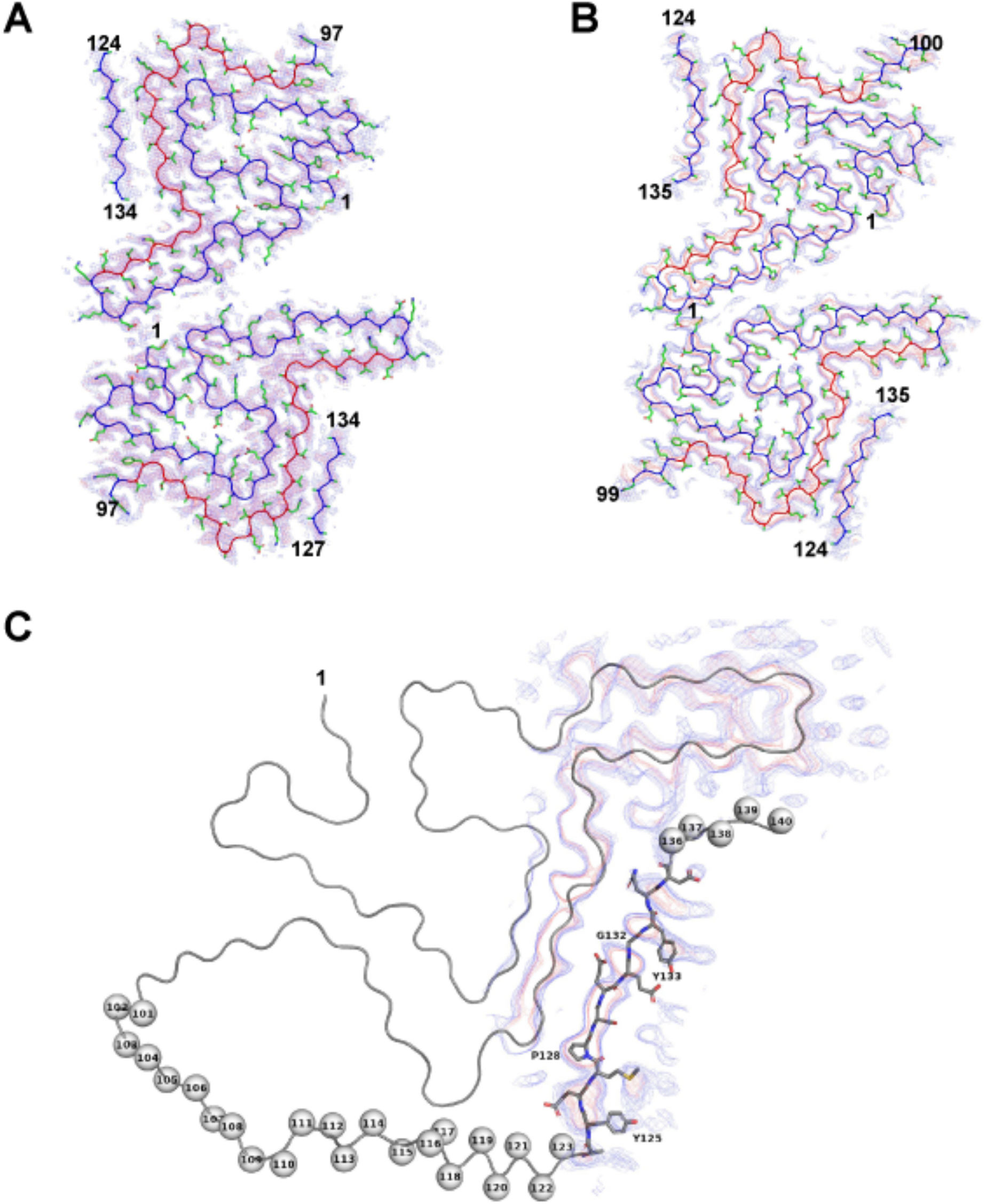
Superimposition of density map and protein backbone for Polymorph 6b and 6c. The backbone was split into two parts (residues 1-60 & 96-100 in blue and the NAC domain (residues 61-95) in red) in individual folds. (**A**) Polymorph 6b backbones fitted in cryo-EM map. (**B**) Polymorph 6c backbone fitted in cryo-EM map at contour of 4 sigma (blue mesh) and 7 sigma (red mesh). (**C**) aSyn amino acid stretch 100-123 unattributed location and distribution. To account for the 60 Å distance from the last visible residue (L100), the density we observe can only be attributed to a sequence spanning residues 118 to 140: DPDN<u>EAY^125^EMPSEEGY^133^QD</u>YEPEA. Y125 and Y133 can be positioned in the two protruding density bulges at both ends of the density. A low-density break in the middle of the strand is compatible with a glycine residue (G132), and a particular density shape is compatible with P128 residue. Map contour in panel A and B: blue mesh (4σ) and green mesh (7σ). In panel C: blue mesh (5σ) and red mesh (7σ).

**Figure 3—figure supplement 1.**
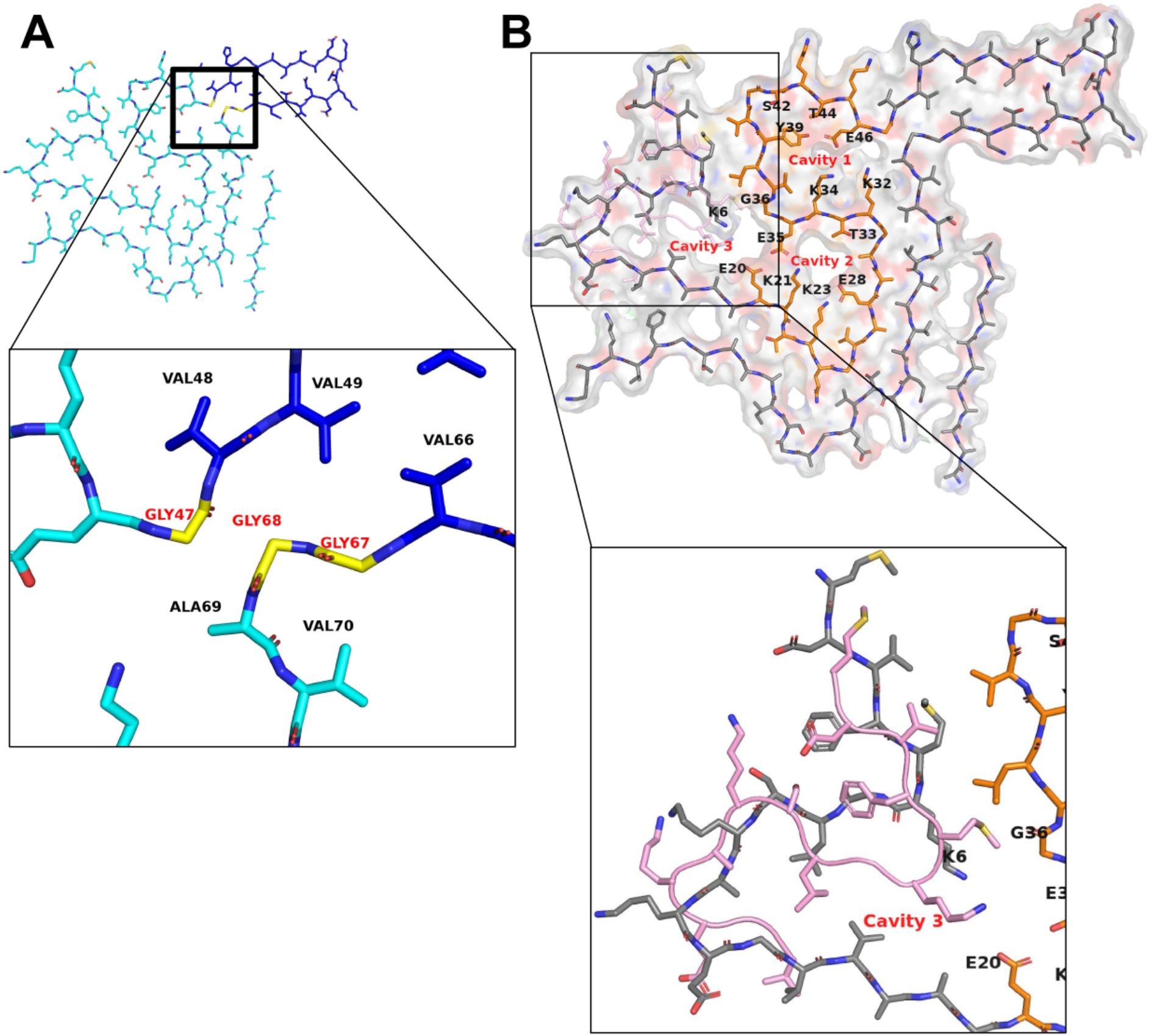
Glycine hinge and cavities in aSyn Polymorph 6 structure. (**A**) Glycine hinge bridging the protruding hairpin (residues 48-66 in dark blue) to N and C-terminal parts (residues 1-46 and 69 to 140, in cyan). Glycine G47, G67-G68 are involved in intermolecular parallel β-sheet hydrogen bonding along the fibril axis through their carbonyl to nitrogen groups. (**B**) aSyn Polymorph 6 structure presents 3 internal hydrophilic cavities buried within the core of the protofilaments and running along the fibrillar axis. The N terminal residues 20 to 47 (in orange) fold into a S shaped structure that creates two hydrophilic holes containing side chains of residues E20 K21, K23, E28 T33, E35 and residues K32, K34, Y39, S42, T44, E46. In Polymorph 6a, the N-terminal residues 6 to 20 form a third hydrophilic hole made of K6, E20, E35 and G36. In the case of Polymorph 6b, residues 1 to 15 (colored in pink) fill the cavity.

**Figure 3—figure supplement 2.**
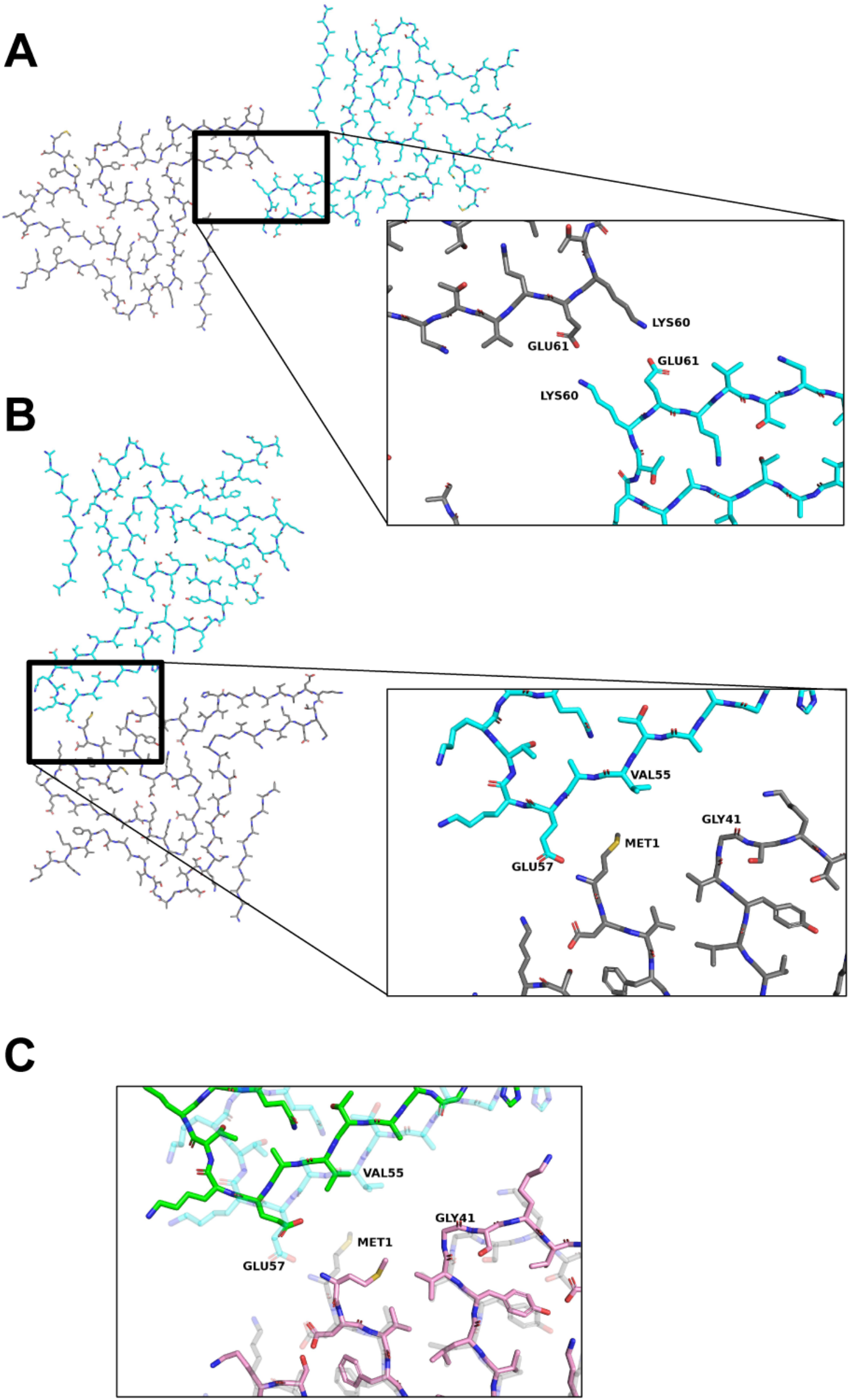
Protofilament interface within aSyn Polymorph 6a, Polymorph 6b and Polymorph 6c. (**A**) The aSyn Polymorph 6a interface is composed of a symmetrical pair of salt bridges formed between the side chains of K60 and E61. (**B**) The asymmetrical Polymorph 6b protofilament interface involves electrostatic interaction between the N terminal M1 amino acid residue and the carbonyl group of amino acid residue E57 side chain. A Van-der-Waals interaction might occur between M1 side chain and residues V55-A56. (**C**) Superposition of Polymorph 6b (cyan and grey chains) and Polymorph 6c (green and pink chains). The inter-protofilament interface within Polymorph 6c is larger than that observed in Polymorph 6b.

**Figure 4—figure supplement 1.**
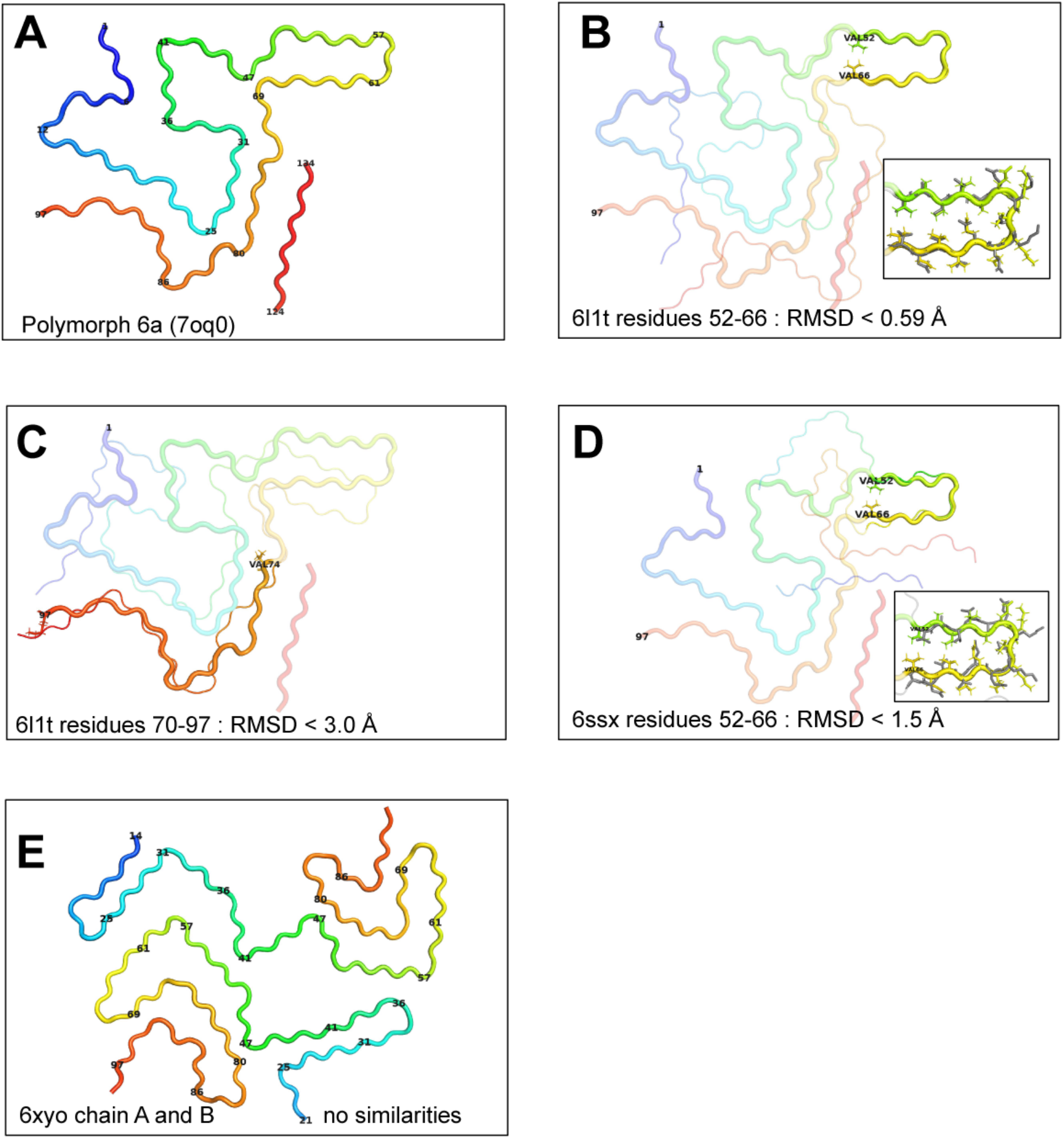
Similarities between Polymorph 6 structure (7oq0) with 6L1T (Y39 phosphorylated aSyn filament), 6ssx (aSyn Polymorph 2a filament) and 6xyo (MSA type I brain extracted filament). (**A**) Polymorph 6a backbone colored as a rainbow from the N (blue) to the C terminal end (red) following the color code in Figure 2A. (**B**) Superimposition of 6L1T (thin cartoon representation) residues 52 to 66 with Polymorph 6. The RMSD value between the two structures is 0.54 Å over CA residues 52 to 66. Inset: all atoms sticks representation of overlapping residues 52 to 66. (**C**) Superimposition of 6L1T (thin cartoon representation) residues 74 to 97 with Polymorph 6a. The RMSD value between the two structures is below 3 Å over CA residues 74 to 97. (**D**) Superimposition of 6ssx (aSyn Polymorph 2a, thin cartoon representation) residues 52 to 66 with Polymorph 6a. The RMSD value between the two structures is below 1.5 Å over CA residues 52 to 66. Inset: all atoms sticks representation of overlapping residues 52 to 66. (**E**) Brain extracted MSA type I filament (6xyo) backbone colored as a rainbow from amino acid residue G14 (light blue) to amino acid residue Q99 (dark orange). No similarity was found with the Polymorph 6 structure.

